# A Neuronal Gene Expression Program Underlying Sustained Network Activity

**DOI:** 10.64898/2026.06.09.731204

**Authors:** J. Wren Kim, Rebecca Eliscu, Adeline J.H. Yong, Sua Lee, Haeun Park, Yuh Nung Jan, Taeyoung Hwang, Liana F. Lareau, Nicholas T. Ingolia

## Abstract

Neurons function within interconnected networks and must continuously adjust their firing properties to maintain stable network activity. Although these adjustments require long-lasting molecular changes, the neuronal gene expression programs that support sustained network activity remain largely unknown. To address this, we developed CalTRAP-seq, a method for profiling gene expression in active neurons via calcium-dependent ribosome tagging. Applying CalTRAP-seq to primary neurons exhibiting synchronous network bursting revealed a gene expression program distinct from stimulus-induced immediate early gene responses, instead enriched for regulators of neuronal excitability. This program was accompanied by widespread alternative splicing of synaptic genes. Notably, neurons participating in network activity exhibited increased nuclear speckle formation, condensates implicated in splicing regulation. Disruption of nuclear speckles impaired synchronous burst dynamics. Together, these findings identify a gene expression program associated with sustained network activity that is complementary to stimulus-responsive gene expression, providing insight into how neurons coordinate gene expression to support stable network function.

## Introduction

Neurons inherently organize into networks, and these networks often exhibit sustained activity without external stimulation. To maintain network stability, these activity levels are tightly regulated by neuronal homeostatic mechanisms that fine-tune synaptic connectivity and intrinsic excitability in response to ongoing network activity^1^. These adjustments require long-lasting molecular changes—such as reorganization of synaptic proteins, ion channels, and other cellular components—which in turn depend on precise regulation of gene expression. We therefore postulate that neurons exhibit dedicated gene expression profiles according to their sustained activity levels, reflecting distinct states of neuronal homeostatic pathways. These profiles then establish distinct cellular phenotypes— referred to here as neuronal ‘activity states’.

However, the distinct gene expression profiles associated with different neuronal activity states have yet to be characterized. A major challenge has been the lack of methods capable of directly linking gene expression patterns with specific activity levels. We previously developed Cal-ID, an engineered biotin ligase that responds to calcium influx by labeling proximal proteins, enabling biochemical enrichment of proteins from highly active neurons^2^. Our work and others^3,4^ established that calcium-dependent protein biotinylation can record neuronal activity in vivo, leading us to hypothesize that it could be adapted for selective gene expression profiling from neurons in highly active state.

In this study, we developed CalTRAP (Calcium-dependent Tagging of Ribosomes and Affinity Purification), a calcium-dependent ribosome tagging technology designed to capture gene expression signatures of active neurons. By applying CalTRAP followed by RNA-seq (CalTRAP-seq) to mouse primary cortical neurons exhibiting spontaneous and synchronous bursting, we uncovered a previously unrecognized gene expression program associated with active neuronal states. This program features extensive regulation of genes controlling synaptic connectivity and intrinsic excitability, including broad changes in splicing patterns of numerous synaptic genes. Intriguingly, we found that the formation of nuclear speckles—membraneless nuclear domains that concentrate splicing factors and serve as hubs for RNA processing^5^— correlates with neuronal activity levels. Our results also showed that disruption of nuclear speckle integrity interferes with synchronous bursts, establishing a functional link between this condensate-mediated RNA regulation and coherent network dynamics. Collectively, our findings reveal a neuronal gene expression program in neurons that is crucial for neuronal network activity, providing new insight into the regulatory mechanisms underlying stable network function.

## Results

### Activity-Dependent Gene Expression and Network Activity

Synchronous network activity is an intrinsic property of neocortical circuits^6^, and this can be modeled in primary neurons cultured in vitro. Dissociated mouse cortical neuron cultures develop robust spontaneous activity, which evolves into coordinated network-wide bursts^7^. We monitored this activity in DIV 26 primary cortical neurons using the calcium indicator jGCaMP8s^8^. Live imaging confirmed that these cultures displayed synchronous bursts of activity across whole networks, with an average duration of 5.86 seconds and a frequency of 4.11 bursts per 3-minute window (Fig. 1a). Notably, when observing the entire culture, these synchronous bursts engaged a large fraction of neurons, while a subset remained inactive. Furthermore, the activity states of individual neurons remained largely stable throughout the monitoring period (Supplementary Video 1).

**Fig. 1.**
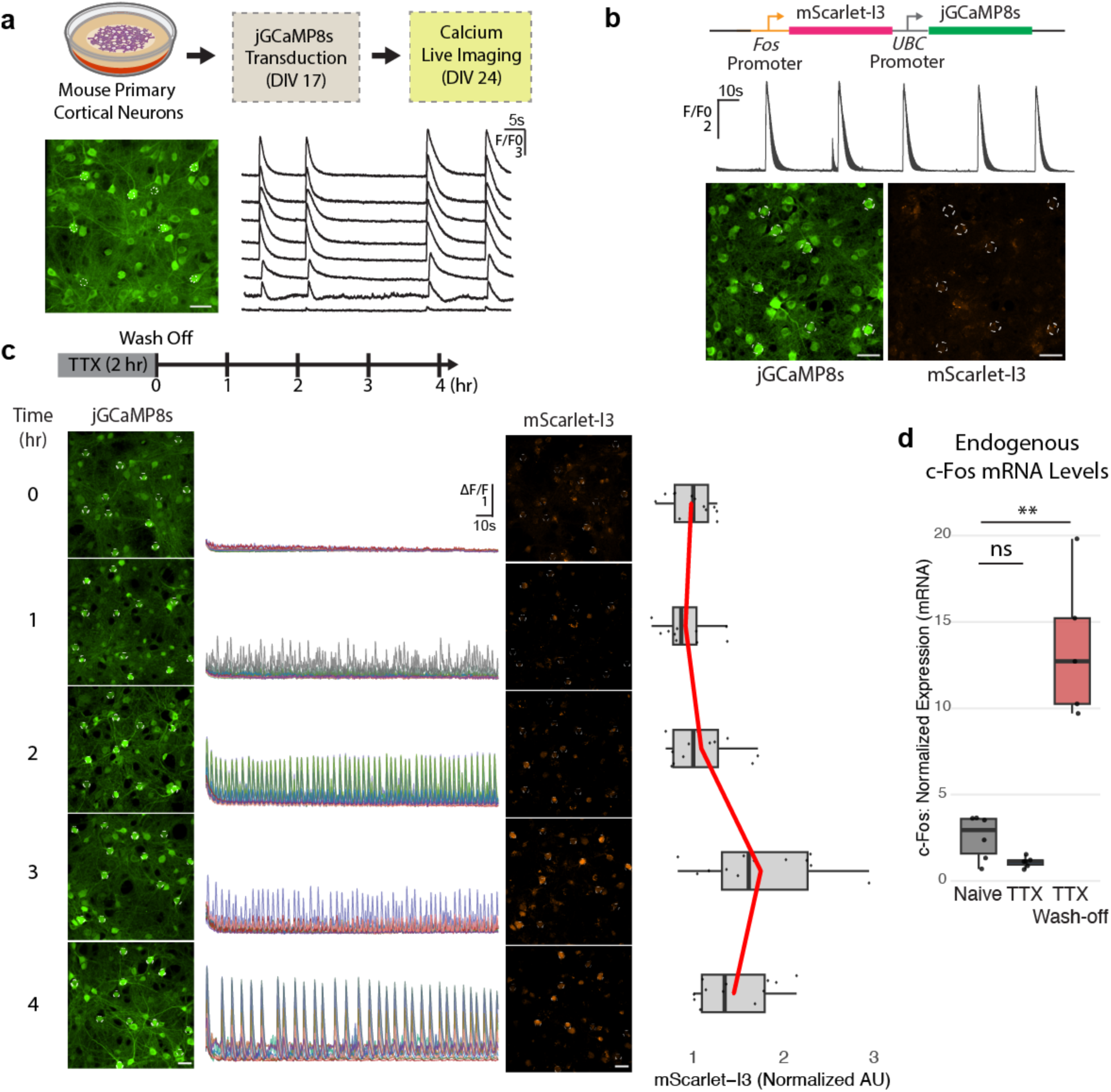
Activity-dependent gene expression and network activity. (**a**) Representative calcium imaging of mouse primary cortical neurons. DIV: days in vitro. **(b)** Calcium and *Fos* promoter dual activity reporter delivered by lentivirus. Calcium traces are from 8 neurons (white circle) showing robust activity, albeit minimal mScarlet-I3 levels. **(c)** Activity suppression using TTX (1 µM) followed by wash-off and activity recovery. Calcium and *Fos* promoter activity (mScarlet-I3 expression) was monitored via live imaging. Representative images of 3 independent trials. (**d**) Endogenous *Fos* mRNA levels measured by qPCR. TTX (1 µM) was treated for 2 hours, total RNA was harvested 1 hour after wash-off. Statistical analysis: one-way ANOVA with Tukey’s post-hoc test. n = 6 each. F = 13.44, p = 0.912 (Naive vs. TTX), p = 0.008 (Naive vs. TTX wash-off). ** p < 0.01. AU: arbitrary unit. Scale bars: 50 µm.

During synchronous bursts, participating neurons fire at high frequencies^9^. Because these neurons experience elevated activity levels during bursts, we asked whether these repeated bursts trigger an activity-dependent gene expression program, characterized by stimulus-triggered induction of immediate early genes (IEGs), such as *Fos*^10,11^. To simultaneously monitor calcium and *Fos* promoter activity, we generated a dual-reporter construct expressing mScarlet-I3^12^ under the control of the *Fos* promoter and jGCaMP8s under the constitutive *UBC* promoter. Encoding both reporters in the same plasmid ensures that reporter expression is analyzed in the same neurons as calcium signals. We found that despite robust synchronous bursts, the *Fos* promoter remained largely uninduced in these active neurons (Fig. 1b).

We wanted to confirm that the reporter could respond to neuronal stimulation. We originally attempted to depolarize the whole culture with 50 mM KCl, but even brief (1-minute) KCl exposure at DIV 24-26 frequently caused substantial neurite degeneration over the following hours, indicating neuronal stress. As an alternative, we silenced network activity with tetrodotoxin (TTX) for 2 hours and then restored it by washing off TTX. This protocol reliably induced *Fos* reporter expression, with the mScarlet-I3 fluorescence peaking 3 hours after the wash-off (Fig. 1c). As protein accumulation lags behind mRNA expression, this indicates that *Fos* induction occurred earlier than 3 hours. Quantitative PCR (qPCR) further confirmed that endogenous *Fos* transcript levels remained low under naïve conditions, dropped to near-minimal levels during TTX treatment, and rapidly increased within 1 hour of network activity recovery following TTX wash-off (Fig. 1d).

Immunostaining in DIV 25 neurons showed that, although levels of phosphorylated CREB— a key marker of calcium signaling and mediator of activity-dependent transcription^13^— were similar between the naïve and TTX wash-off conditions, endogenous c-Fos protein levels were significantly lower in the naïve condition (Extended Data Figs. 1a to 1c).

Together, these results demonstrate that although neurons engaged in network activity exhibit robust calcium transients and associated signaling such as CREB phosphorylation, this activity does not necessarily trigger *Fos* promoter induction.

### Calcium-Dependent Tagging of Ribosomes and Affinity Purification (CalTRAP)

The absence of stimulus-responsive gene induction in neurons exhibiting robust and continuous firing led us to hypothesize the existence of a distinct gene expression program defining these neuronal activity states. Identifying these programs requires a method to selectively profile gene expression in neurons engaged in network activity. To achieve this, we adapted Cal-ID, the calcium-dependent protein labeling enzyme that we previously developed as a biochemical recorder of neuronal activity^2^. Cal-ID achieves calcium- and thus activity-dependent protein biotinylation using a circularly permuted TurboID^14^ with calmodulin-dependent intramolecular complementation. We further developed this strategy into a calcium-dependent ribosome labeling system, termed CalTRAP (Calcium-dependent Tagging of Ribosomes and Affinity Purification). CalTRAP consists of two key components: 1) CalBirA, a calcium-dependent and substrate-specific biotin ligase, or its calcium-independent control, circularly permuted BirA (cpBirA); and 2) the ribosomal protein uL1 (Rpl10a) fused to the substrate, an AviTag biotin acceptor sequence (Figs. 2a and 2b). In this system, an influx of calcium triggers biotinylation of ribosomes, enabling the selective capture of ribosomes and their associated mRNAs from neurons during network activity (Figs. 2c and 2d).

**Fig. 2.**
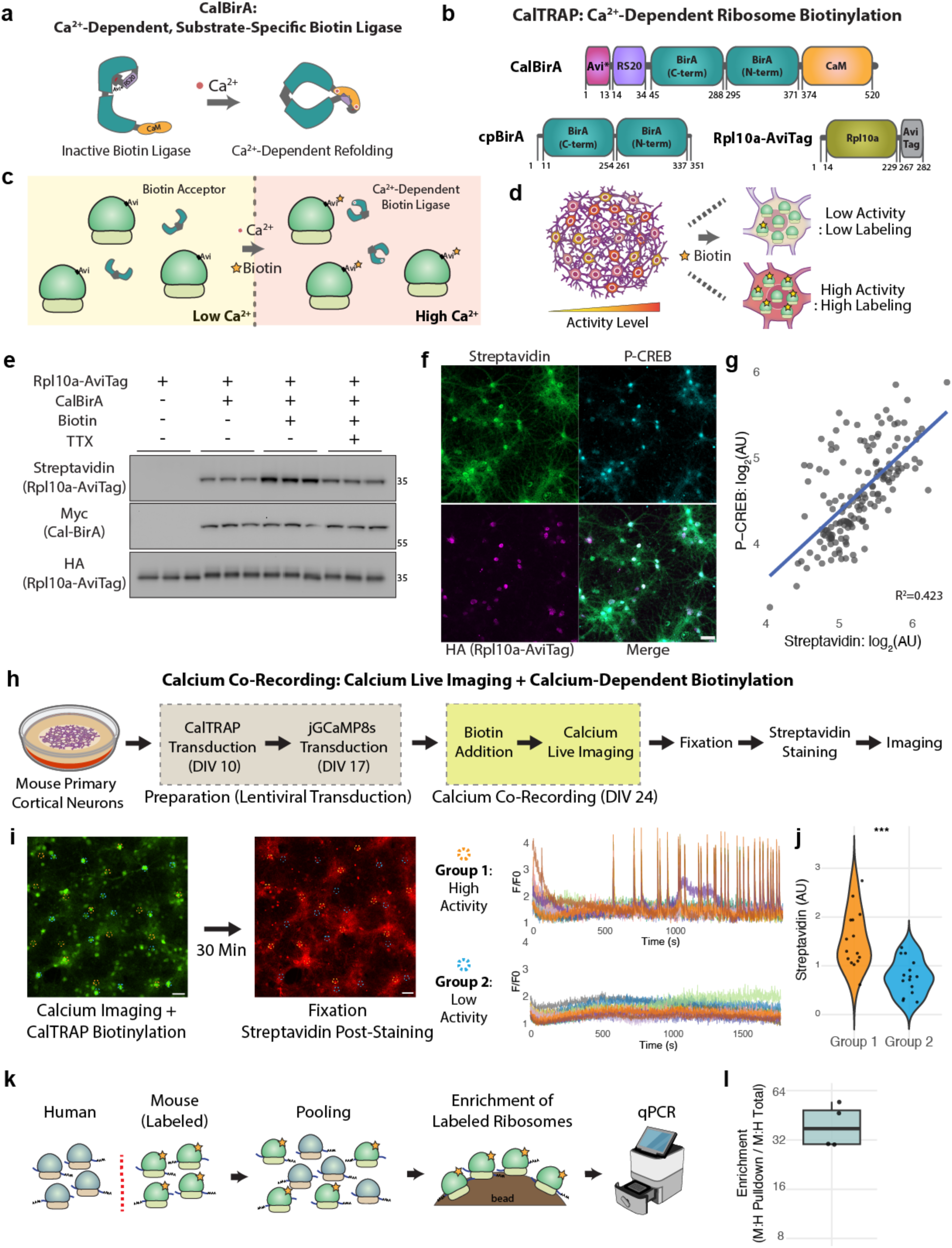
CalTRAP enables selective ribosome labeling from active neurons. (**a**) A schematic diagram of a calcium-dependent, substrate-specific biotin ligase via calcium-dependent refolding. (**b**) CalTRAP system for calcium-dependent ribosome labeling. (**c** and **d**) CalTRAP for selective gene expression profiling from active neurons. (**e**) Activity-dependent ribosome biotinylation. Mouse primary cortical neurons at DIV 19. Representative blot (n = 4). Biotin: 100 µM, TTX: 1 µM. TTX was pre-incubated for 1 hour before 30 mins of biotinylation. Numbers on the right: kDa. (**f** and **g**) Correlation between CalTRAP biotinylation and calcium signaling. Neuron at DIV 22. n = 162. Scale bar: 50 µm. (**h** to **j**) Calcium co-recording with simultaneous calcium live imaging and CalTRAP biotinylation. n = 15 per group. Welch’s t-test, p < 0.0003. Scale bar: 100 µm. (**k** and **l**) CalTRAP enrichment test using pooled lysates. n = 4.

We first validated that CalTRAP biotinylation is coupled to neuronal activity. In primary cortical neurons expressing CalTRAP, western blot analysis revealed robust biotinylation of Rpl10a-AviTag upon biotin addition (Fig. 2e). This biotinylation was reduced by treatment with TTX, which blocks neuronal firing. Furthermore, immunocytochemistry showed a robust positive correlation between CalTRAP biotinylation levels and p-CREB (Figs. 2f and 2g). To directly correlate activity with CalTRAP biotinylation, we performed live calcium imaging alongside CalTRAP labeling. We co-expressed jGCaMP8s and CalTRAP in neurons, added biotin, and recorded calcium activity via live imaging for 30 minutes before fixing and staining with fluorescent streptavidin (Fig. 2h). Neurons engaged in ongoing network activity showed significantly higher biotinylation compared to silent neighbors (Figs. 2i and 2j, Supplementary Video 2).

Different neuronal types can exhibit distinct baseline activity levels. These baseline differences have the potential to influence CalTRAP labeling efficiency and introduce bias into gene expression profiles. To test this, we labeled neurons with cpBirA or CalBirA and compared biotinylation levels between excitatory (Camk2a-positive) and inhibitory (Gad67-positive) neurons using immunocytochemistry. Both calcium-independent and calcium-dependent labeling yielded comparable biotinylation across neuronal types (Extended Data Figs. 2a to 2d). Of note, inhibitory neurons showed slightly higher labeling under both conditions, likely driven by cell type-related differences in promoter activity and enzyme expression levels.

Finally, we confirmed the selectivity of biotinylated ribosome pulldown. We expressed CalTRAP in mouse cortical neurons and pooled their lysates with lysates from HEK293T cells, which lack CalTRAP (Fig. 2k). After enriching biotinylated ribosomes with streptavidin beads under stringent high-salt washes^15,16^, we purified ribosome-bound mRNAs. qPCR analysis demonstrated a greater than 30-fold enrichment of mouse neuron-specific transcripts (*Map1a*) over HEK293T-specific transcripts encoding SV40 Large T antigen (Fig. 2l). Collectively, these results establish CalTRAP as a robust tool for selective gene expression profiling from neurons in high activity states.

### Network Activity and Genes Regulating Neuronal Excitability

To profile neuronal gene expression associated with network activity, we performed CalTRAP-seq with DIV 24 neurons under three conditions: (1) Control TRAP, using calcium-independent cpBirA as a baseline that labels all neurons regardless of activity level; (2) CalTRAP-SR, serving as a positive control for stimulus-responsive gene expression induced by network activity silencing and recovery; and (3) CalTRAP-NA, which captures gene expression from neurons with native network activity (Figs. 3a and 3b). Biological replicates showed high correlation with each other (Extended Data Figs. 3a to 3c); in total, 13781 genes were analyzed for differential expression^17^ after filtering (Supplementary Table 1).

**Fig. 3.**
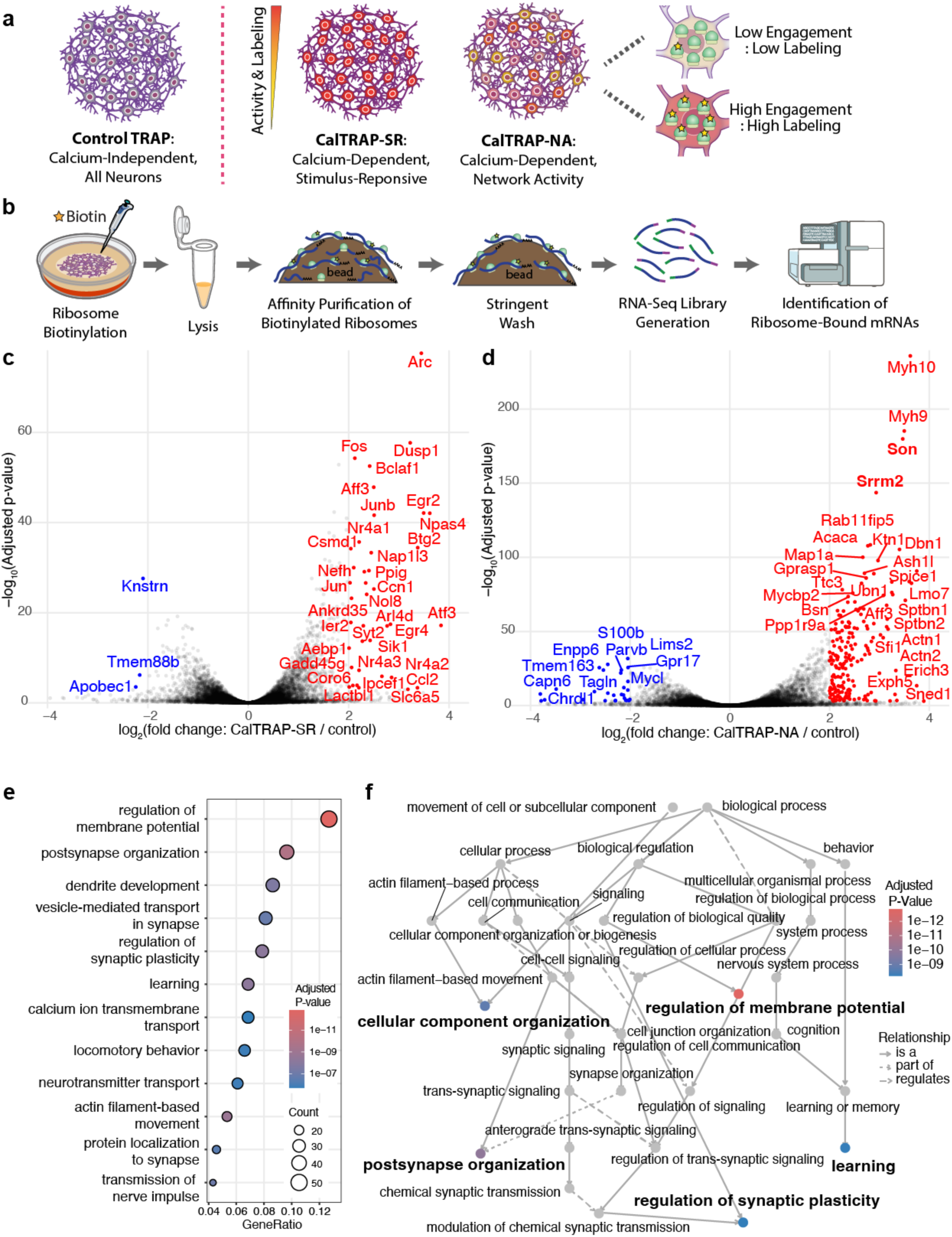
Profiling gene expression associated with spontaneous network activity. (**a** and **b**) CalTRAP-seq experimental design with primary mouse cortical neurons. Control TRAP: calcium-independent cpBirA. CalTRAP-SR condition: TTX (1 µM) for 2 hours followed by wash off. All conditions: 1 hour labeling with exogenous biotin addition (100 µM). Biological replicates (n = 2) per condition. (**c** and **d**) Volcano plots comparing differential gene expression between (c) CalTRAP-SR vs. Control TRAP or (d) CalTRAP-NA vs. Control TRAP. Genes with adjusted p-value < 0.001 and log2FoldChange > 2 or < −2 are highlighted. (**e** and **f**) Gene ontology analysis (biological processes) with the significantly enriched genes from (d) against the total 13781 CalTRAP-seq detected genes reveals pathways regulating neuronal excitability.

Comparing CalTRAP-SR to Control TRAP revealed 39 genes with highly significant enrichment (adjusted p-value < 0.001 and log_2_ fold change > 2) (Fig. 3c). As expected, stimulation activates a classic stimulus-responsive gene program, marked by the robust induction of immediate-early genes—such as *Arc*, *Fos*, *Dusp1*, *Egr2*, and *Npas4*—in agreement with well-established literature^18^. In contrast, the comparison of CalTRAP-NA to Control TRAP identified a distinct set of 199 highly enriched genes (adjusted p-value < 0.001 and log_2_ fold change > 2) specific to neurons engaged in ongoing network activity (Fig. 3d). A direct comparison between CalTRAP-SR and CalTRAP-NA gene expression profiles further demonstrates that the gene expression program of native network activity is distinct from the canonical stimulus-response program (Extended Data Fig. 4a).

We noticed that the most significantly enriched genes from the CalTRAP-NA condition, such as *Myh10*, *Rab11fip5*, *Dbn1*, and *Map1a*, have been implicated in synaptic regulation and neurodevelopment^19–22^. To gain insight into the biological processes associated with network activity–associated genes, we performed gene ontology (GO) analysis on genes significantly upregulated in CalTRAP-NA relative to control TRAP. GO analysis revealed enrichment for processes related to neuronal excitability, including membrane potential regulation and synaptic function (Figs. 3e and 3f). Using SynGO annotations^23^, we further asked whether the most enriched genes are associated with synaptic function. Notably, 57 of the 199 enriched genes are annotated as synaptic genes in SynGO (Extended Data Fig. 4b, Supplementary Table 1).

To further test potential cell-type bias, we analyzed snRNA-seq data from cortical neuronal cultures at DIV 23^24^. We examined expression of 15782 genes from 2365 cells and identified the most significant excitatory-and inhibitory-specific genes (37 and 23 genes, respectively, Supplementary Table 1). These cell type–specific genes did not show biased expression in CalTRAP pulldown results (Extended Data Figs. 5a and 5b), supporting that CalTRAP does not introduce significant cell type–dependent bias in gene expression profiling. Together, these results from CalTRAP-seq reveal a previously uncharacterized gene expression program in neurons engaged in network activity. Notably, this program is enriched for regulators of intrinsic excitability and synaptic connections, highlighting its potential role in maintaining stable and coherent network activity.

### Network Activity and Nuclear Speckle Formation

Our CalTRAP-seq analysis revealed significantly increased expression of *Srrm2* and *Son* genes in neurons engaged in network activity (Fig. 3d). These genes encode core scaffolding proteins of nuclear speckles, membraneless nuclear condensates that serve as key mRNA processing hubs. We hypothesized that neuronal activity states are linked to speckle formation (Fig. 4a). Immunocytochemistry confirmed a positive correlation between nuclear speckle abundance and levels of p-CREB (Figs. 4b and 4c). We directly visualized nuclear speckles in active neuronal networks using a fluorescent reporter in conjunction with live calcium imaging in mouse cortical neurons. We expressed the calcium indicator jGCaMP8s by lentiviral transduction and delivered a full-length human SRRM2-mScarlet-I fusion protein under the human *EF1α* promoter via nucleofection due to its large size. Live imaging was performed at DIV 26, and we compared SRRM2-mScarlet-I signals between neurons engaged in synchronous bursts and non-participating neurons (Fig. 4d). The results revealed that neurons involved in synchronous network activity exhibited significantly higher levels of nuclear speckle formation (Figs. 4e and 4f).

**Fig. 4.**
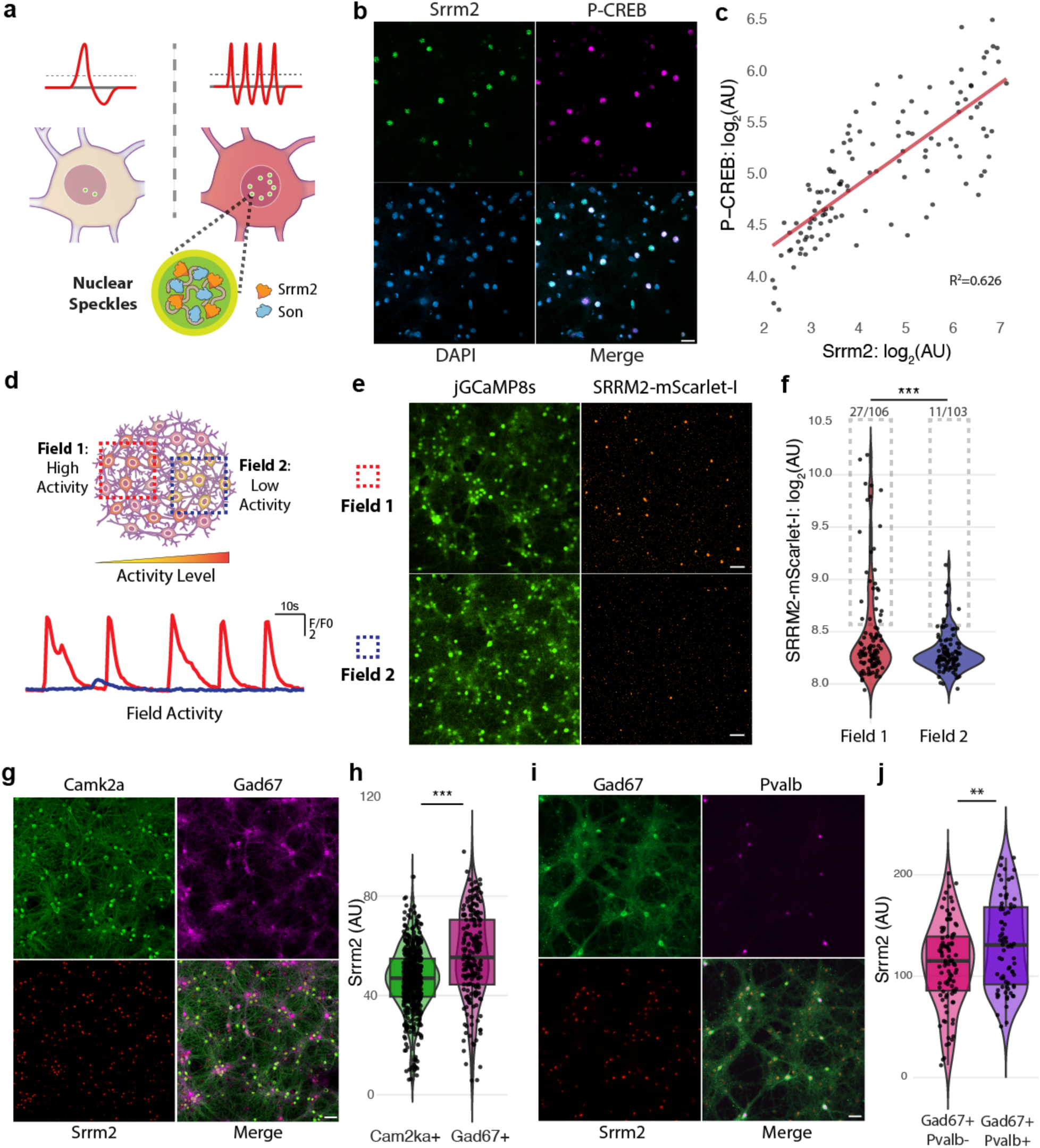
Nuclear speckle formation in neurons engaged in network activity. (**a**) Relationship between nuclear speckles and neuronal activity. (**b** and **c**) Correlation between P-CREB and nuclear speckles. Primary mouse cortical neurons at DIV 23. n = 119. Scale bar: 50 µm. (**d** to **f**) Nuclear speckle was monitored between high and low network activity areas in the same culture. Dotted box visualizes the 4^th^ quantile. Welch’s t-test, n = 106, p = 0.00027. Scale bar: 50 µm. (**g** and **h**) Nuclear speckle formation between excitatory (Cam2ka+) and inhibitory (Gad67+) neurons. Primary cortical neurons at DIV 24. n = 507 (Camk2a+), n = 203 (Gad67+). Welch’s t-test, p = 6.258e-11. Scale bar: 50 µm. (**i** and **j**) Nuclear speckle formation between Pvalb- vs. Pvalb+ inhibitory neurons. Nuclear speckle formation between excitatory (Cam2ka+) and inhibitory (Gad67+) neurons. Primary cortical neurons at DIV 23. n = 97 (Gad67+/Pvalb-), n = 75 (Gad67+/Pvalb+). Welch’s t-test, p = 0.003788. Scale bar: 50 µm.

To address potential cell-type-specific differences in nuclear speckle formation related to varying baseline activity levels, we co-stained neurons with excitatory (Camk2a) and inhibitory neuronal markers (Gad67). The results demonstrated that inhibitory neurons form nuclear speckles more frequently than their excitatory counterparts (Figs. 4g and 4h). This difference may be linked to the higher baseline firing rates of inhibitory neurons. In this regard, we compared nuclear speckle formation between parvalbumin-positive (PV+) neurons and other inhibitory neurons, since PV+ interneurons are known to exhibit characteristically high baseline firing rates and serve as primary regulators of network activity^25^. We found that PV+ neurons (which are both Gad67-and Pvalb-positive) exhibit significantly higher levels of nuclear speckle formation than other inhibitory neurons (which are Gad67-positive but Pvalb-negative) (Figs. 4i and 4j). Together, these data demonstrate that the formation of nuclear speckles is correlated with neuronal network activity levels. In addition, high levels of speckle formation are observed in PV+ neurons, consistent with their distinctively high baseline activity.

### Network Activity and Alternative Splicing of Synaptic Genes

Given the established role of nuclear speckles in coordinating alternative splicing, we next asked whether neurons engaged in network activity exhibit distinct splicing patterns. We analyzed our CalTRAP-seq data using rMATS^26^ to compare splicing patterns between the CalTRAP-NA and control TRAP conditions. This revealed a total of 1,554 alternative splicing events with significant different usage (FDR < 1×10⁻⁵), predominantly exon inclusion and exclusion events (76.2% of all events) (Fig. 5a, Supplementary Table 2).

**Fig. 5.**
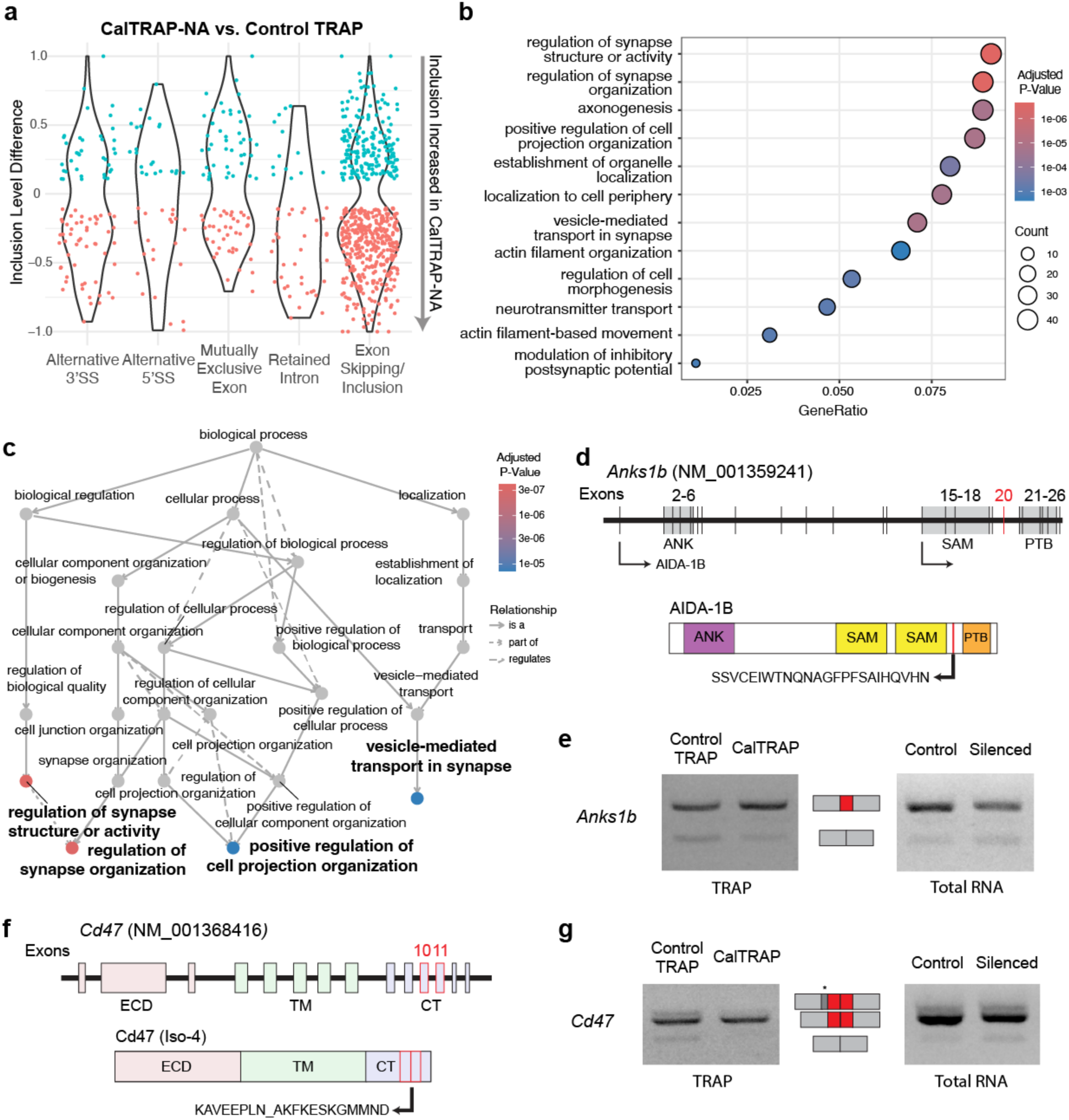
Network activity-associated alternative splicing of synaptic genes. (**a**) Alternative splicing analysis between CalTRAP-NA and control TRAP shows predominant exon skipping/inclusion events. (**b** and **c**) Gene ontology: biological processes with alternatively spliced genes reveals active splicing regulation of synaptic genes. Total 458 genes were selected from significant exon skipping/inclusion events (FDR < 1e-05). Reference universe is the 13781 genes detected from CalTRAP-seq. (**d** to g) (d and e) *Anks1b* exon 20 (f and g) *Cd47* exons 10 and 11 show elevated inclusion from CalTRAP pulldown and reduced inclusion by silencing. Silencing: TTX (1 µM), NBQX (10 µM), AP5 (1 µM) for 24 hours. *Exon not included in the NM_001368416 annotation: exon 9 of XR_384529. Representative images from 4 pulldown experiments, 8 silencing experiments. ANK: Ankyrin repeats, SAM: sterile alpha motif, PTB: phosphotyrosine binding, ECD: extracellular domain, TM: transmembrane, CT: C-terminal domain. Primary mouse cortical neurons at DIV 21.

Gene ontology analysis of genes with significant differences in alternative splicing demonstrated that these genes are predominantly synaptic (Figs. 5b and 5c), indicating that the network activity–associated gene expression program regulates not only transcript abundance but also isoform selection of synaptic genes. Of note, we found that network activity–associated splicing regulation led to complex changes in transcript isoform diversity, with 75 genes identified with more than two splicing events each (Supplementary Table 2). These include genes previously known to be alternatively spliced in an activity-dependent manner. Our analysis detected network activity–associated changes in three known alternative splicing events in the gene *Nrxn1*, which encodes the presynaptic protein neurexin-1 (Extended Data Fig. 6a)^27,28^. Our analysis also detected significant differences in the splicing patterns of microexons in *Eif4g1* and *Eif4g3*, which encode translation initiation factors eIF4G1 and eIF4G3, in the CalTRAP-NA condition (Extended Data Fig. 6b). Microexon inclusion in these genes is known to be regulated by neuronal activity and is important for synaptic local translation^29^.

Our results uncovered previously unknown associations between network activity and alternative splicing of key synaptic genes. *Anks1b*, which encodes the postsynaptic scaffold protein AIDA-1 and is linked to various psychiatric disorders^30^, showed network activity–associated splicing of exon 20. This exon encodes a part of the protein adjacent to the phosphotyrosine binding (PTB) domain, which is critical for interactions with other synaptic proteins^31,32^ (Fig. 5d). PCR amplicons from CalTRAP pulldown cDNA confirmed that exon 20 inclusion is elevated in neurons engaged in network activity (Fig. 5e).

Furthermore, Cd47, a membrane protein that serves as a "don’t eat me" signal to prevent microglial phagocytosis during synaptic pruning^33^, exhibited network activity–associated alternative splicing of exons 10 and 11(Figs. 5f and 5g). These exons encode the cytoplasmic tail, a region important for cellular signaling and protein turnover^34^.

We then tested whether silencing neuronal activity would influence splicing of these exons. Treatment with TTX along with AP-5 and NBQX (which inhibit NMDARs and AMPARs, respectively) for 24 hours reduced inclusion of the target exons in *Anks1b* and *Cd47* (Figs. 5e and 5g), showing that manipulation of sustained activity states leads to alternative isoform choices of these genes. Collectively, these results demonstrate that neurons engaged in network activity employ a distinct alternative splicing program to fine-tune the proteoform expression of key synaptic genes and regulators. These findings further suggest a sophisticated mechanistic effect from neuronal activity states on post-transcriptional regulation in supporting stable network function.

### Perturbation of Nuclear Speckles Impairs Network Activity

The tight coupling between neuronal activity state and alternative splicing of synaptic genes led us to hypothesize that nuclear speckles are functionally important for generating and maintaining synchronous network dynamics. To test this, we used CRISPR interference (CRISPRi) to knock down *Srrm2* and thereby disrupt nuclear speckle integrity. We designed guide RNAs that reduced *Srrm2* mRNA expression by more than 75% when delivered via lentivirus at DIV 14, a time point chosen to minimize impacts on early neuronal development (Extended Data Fig. 7a). We then monitored network activity using calcium imaging at DIV 25. This perturbation resulted in aberrant network activity patterns.

Specifically, *Srrm2* knockdown led to overall elongated burst durations without apparent changes in burst amplitude or the total number of bursts (Figs. 6b and 6c, Extended Data Figs. 7b and 7c, Supplementary Video 3 and 4). These results demonstrate that the structural and functional integrity of nuclear speckles, maintained by Srrm2, is critically required for the precise coordination of synchronous network activity.

**Fig. 6.**
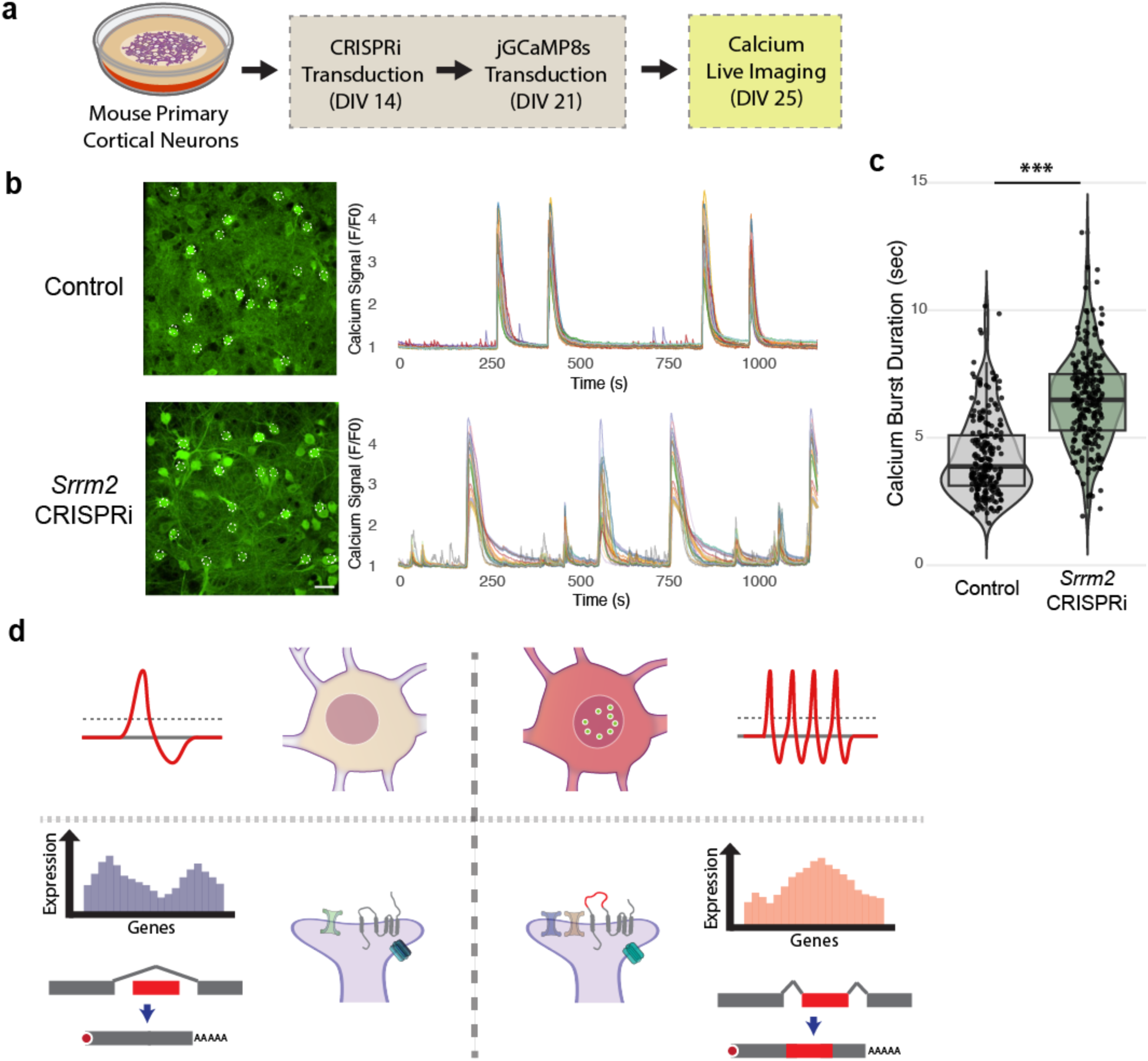
Srrm2 CRISPRi disrupts the integrity of network activity. (**a** to **c**) CRISPR interference of Srrm2 increases calcium burst durations. (b) Representative traces from 20 neurons. (c) Traces: n = 210 from 11 biological replicates (control), n = 251 from 13 biological replicates (CRISPRi). Welch’s t-test, p < 2.2e-16. Scale bar: 50 µm. (**d**) Differential gene expression and alternative splicing synaptic genes shape activity-dependent cellular states in neurons.

## Discussion

The coupling between gene expression and electrophysiological activity is fundamental to the homeostatic regulation of brain function, and its disruption is a hallmark of many neurodevelopmental and neurodegenerative disorders. Historically, activity-dependent gene expression has been studied primarily in response to discrete stimuli, leading to seminal discoveries about the mechanisms of learning and memory. However, the gene expression programs that underlie continuous and basal network activity remained largely unknown. This knowledge gap might be, in part, due to a lack of tools for profiling gene expression based on activity level within a heterogeneous neuronal population. In this study, we introduce CalTRAP-seq, a method that couples calcium levels to ribosome tagging, enabling the selective isolation of transcripts being translated from active neurons. By applying CalTRAP-seq to in vitro cortical cultures showing spontaneous synchronous bursts, our study uncovers a previously uncharacterized gene expression program important for maintaining the integrity of neuronal network activity. These findings extend our understanding of activity-dependent gene regulation beyond transient, stimulus-driven responses to include sustained network activity–associated programs. Our results further suggest that neurons with different levels of continuous activity can be viewed as occupying distinct cellular states, each defined by unique gene expression patterns including synaptic proteome and proteoforms (Fig. 6d). This perspective offers a valuable framework for interpreting neuronal gene expression profiles.

Our application of CalTRAP-seq identified the regulation of neuronal excitability as a central theme in the gene expression profile of neurons engaged in network activity. This finding aligns well with requirement for neuronal homeostatic feedback mechanisms fine-tuning synaptic connectivity and intrinsic excitability to establish and maintain coordinated firing across a network^35^. The distinct gene expression patterns we identified thus likely reflect the cellular and molecular processes that enable individual neurons to calibrate and sustain specific activity levels within the network. Importantly, these network activity–associated genes are directly relevant to human disease. Among the most significantly enriched 199 genes in CalTRAP-NA (log2 fold change > 2 and adjusted p-value < 0.001), 38 are classified as high-confidence risk factors (SFARI Category I or II) for autism spectrum disorders (ASD), a number far exceeding that expected by chance (p < 1.2×10^-9^) (Extended Data Fig. 4c, Supplementary Table 1). The convergence of our network activity–associated gene profile with ASD-associated genes suggests that dysregulation of this precisely-tuned regulatory program may contribute to the circuit-level pathophysiology of neurodevelopmental disorders. Thus, understanding the activity-dependent control mechanisms of these genes will provide crucial insights into the mechanisms underlying associated conditions.

Diversifying synaptic proteoforms is a key mechanism for synaptic development and regulation^36^. Our results suggest that activity state-associated nuclear speckle reorganization and alternative splicing of synaptic genes constitute a central mechanism for achieving this diversification. Mutations in nuclear speckle components are implicated in a wide spectrum of brain disorders. For example, mutations in the genes encoding the key scaffolding proteins Srrm2 and Son lead to neurodevelopmental or intellectual disorders^37–39^. However, their regulation and function in the brain remain active areas of inquiry^5,40^. We reveal that neuronal activity levels are a key determinant of these nuclear condensates. Crucially, our experiments demonstrate that disrupting speckle integrity through Srrm2 knockdown impairs synchronous network activity, establishing a critical role for nuclear speckles in maintaining coherent circuit dynamics.

This activity-associated regulation of nuclear speckles has functional consequences: the alternative splicing of a suite of synaptic genes and regulators. We show that *Anks1b* and *Cd47* transcripts are alternatively spliced in neurons engaged in network activity. Although both genes are known to produce multiple isoforms, how these specific splicing events are regulated in the context of neuronal function has remained unclear. The 72-nucleotide exon 20 of *Anks1b* encodes a region flanked by sterile alpha motif (SAM) and PTB domains, both critical for protein localization and binding interactions. Its inclusion varies across AIDA-1 proteoforms, suggesting a functional role for this splicing event^31^. Similarly, inclusion of exons 10 and 11 in *Cd47* varies across isoforms, and these exons contain a TRAF2-dependent ubiquitination site that affects Cd47 protein stability^41^. In addition, our results indicate that the high baseline inclusion of microexons in *Eif4g1* and *Eif4g3*—which is required for their subsequent stimulus-dependent reduction^29^—could be driven by network activity. This observation suggests that certain alternative splicing events previously attributed to acute stimulation may, in fact, be tuned to the neuron’s ongoing activity state. Collectively, these findings reveal sophisticated and likely bi-directional coupling by which different levels of neuronal activity shape synaptic proteoform expression to fine-tune network function.

Since ribosome-bound mRNAs are mature and likely to be under active translation, ribosome pulldown technologies^42,43^ provide significant advantages to investigate splice isoforms leading to distinct proteoforms^44^. While it is based on the principle of proximity labeling, one of key advantages of CalTRAP is its use of abundant ribosomal proteins and cytosolic enzymes, which enables mesoscale ribosome labeling for a comprehensive survey of cellular mRNA translation. Nevertheless, CalTRAP labeling efficiency may still be influenced by distance from the calcium source, and this necessitates careful consideration of mRNA localization when interpreting CalTRAP-seq datasets.

Our findings open several new avenues for investigation. Future work should determine whether distinct neuronal cell types employ unique activity state–dependent gene programs and how different patterns of network activity are reflected in the gene expression profile. These questions are of particular importance in the native and diverse circuit contexts in the brain. In addition, a recent study reported that neuronal stimulation induces nuclear speckle formation in the brain^45^. Elucidating the temporal dynamics of nuclear speckles and alternative splicing regulation in vivo^46^, together with potential protein turnover regulation, will be an important area for future research. Additionally, a major outstanding question is the identity of the specific signaling cascades that transduce neuronal activity states into these gene expression programs to enable feedback regulation, thereby maintaining the homeostatic integrity of neuronal network^47^. Addressing these questions will be essential for a complete understanding of how neurons dynamically regulate their cellular composition and processes to support the complex function of the brain.

## Methods

All animal protocols are in accordance with the regulations of University of California,

Berkeley Animal Care and Use Committee and the National Institutes of Health (NIH) *Guide for the Care and Use of Laboratory Animals.* Animals were housed in a 12-hour reverse dark/light cycle with free access to water and food (LabDiet 5001), at 22.5 °C ambient temperature and with 50 % humidity. All animal procedures were conducted according to protocols approved by the Institutional Animal Care and Use Committee (IACUC) and the Office of Laboratory Animal Care (OLAC) at the University of California, Berkeley (AUP-2018-08-11380).

### Cloning

PCR primers were designed to have overhangs, PCR reactions were performed with Q5 polymerase (NEB), and the fragments were assembled using HiFi (NEB) or Gibson assembly (NEB). Ligated plasmid products were introduced by heat shock transformation into competent Stbl3 (Invitrogen).

### Primary Cortical Neuron Culture

Dissociated primary neurons were prepared from developing brain at embryonic day 18 (CD-1, Charles River). Embryonic cortical tissues were dissected in dissecting medium (Dulbecco’s Modified Eagle Medium (DMEM) with 20 % FBS, 0.5 mM GlutaMax, 6 μM glucose, Gibco), digested with 20 mg/ml papain (Worthington) for 20 min at 37 °C, and plated at a concentration of 2.4 × 10^6^ cells for a 12-well plate (cortical neurons) or 70,000 – 80,000 neurons per one well of 24-well plate (hippocampal neurons), 4 × 10^6^ cells per 100 mm dish. Culture plates were pre-coated with 50 μg/mL poly-D-lysine. Cultures were maintained under Neurobasal Plus (Gibco) medium with a serum-free supplement B-27 plus (Gibco) and 0.5mM GlutaMax (Gibco). After 7 days, culture media was switched to Neurobasal + B-27 (Gibco). All animal procedures were conducted according to protocols approved by the Institutional Animal Care and Use Committee (IACUC) and the Office of

Laboratory Animal Care (OLAC) at the University of California, Berkeley (AUP-2018-08-11380-2).

### Immunocytochemistry

Cells were washed 3 times and fixed with 4 % paraformaldehyde for 15 min at RT, washed 3 times again then permeabilized and blocked for 1 hour with 5 % BSA and 0.1 % saponin in PBS. The blocked cells were subsequently incubated with primary antibody overnight at 4 °C. On the following day, the cells were washed 3 times and incubated with secondary antibody for 1 hour at room temperature in a light controlled condition. After 3 × wash with PBS buffer, the cells were mounted on cover slides with mounting media containing DAPI (VECTASHIELD® Plus, Vector Labs, H-1900). Images were taken with LSM900 (Carl Zeiss) confocal laser scanning microscope under 20 × air or 63 × oil objectives.

### Immunoblotting

Cells were lysed with lysis buffer (20 mM Tris-HCl (pH 7.4), 150 mM NaCl, 1 mM EGTA, 1 % Triton X-100, protease inhibitors) incubated on ice for 10 minutes at 4 °C, and debris was separated by centrifugation for 10 min × 19,600 g at 4 °C. Supernatant was collected, protein concentration was measured, and the lysate was mixed with 2 × Laemmli sample buffer. SDS-PAGE and transfer were performed on Invitrogen Bolt™ system with Bis-Tris 4-12% gradient gels. ProteinSimple blotting system was used for visualization.

### Lentivirus generation

Lentivirus transfer vectors were co-transfected into low passage number (< 15) HEK293T cells with VSV-G and gag/pol plasmids at a 1:2:3 ratio. 1 day after transfection culture, media was replaced with new media with 10 mM (final) HEPES. The first harvest of packaged virus was made 2 days after transfection, mixed with Lenti-X concentrator (Takara), and stored at 4 °C. A second harvest was made 3 days after transfection, mixed with Lenti-X concentrator and concentrated along with the first harvest following the manufacturer’s recommendation. The relative titer of the lentiviral concentrate was estimated using Lenti-X GoStix (Takara).

### Calcium live imaging

Calcium live imaging was performed on a Nikon Eclipse Ti microscope equipped with a Yokogawa CSU-X1 spinning disk confocal unit and 4× or 20× objective lens. Neurons expressing jGCaMP8s were maintained in Neurobasal + B-27 media at 37 °C and 5 % CO2, imaging was performed without switching culture media. Time-lapse fluorescence images were acquired using 488/568 nm excitation controlled with NIS-Elements.

### Image Analysis

For all quantitative image analyses, ImageJ (1.53f51)/FIJI was used^48^. Western blot quantification was conducted with ‘Gels’ analysis function of ImageJ. For the analysis of immunostaining from mouse primary neurons, all neurons in the field were selected via Threshold and Analyze Particles tools with 50 – infinity size with circularity 0.15 – 1 selection in ImageJ. When a subset of neurons was selected, ImageJ plugin ‘ROI 1-click tool’ was used with the appropriate radius to target only soma regions. For the analysis of calcium live imaging, ImageJ plugin ‘ROI 1-click tool’ was utilized to select cell bodies, with radius 1-5 targeting soma region and avoiding the nucleus. Target mean intensity values were extracted from the selections and transferred to R. The results were analyzed in R and visualized using ggplot2 (3.0.0)^49^. Blinding: the experimental conditions were blinded during the image analysis. For burst detection, 1 + 25 % of the maximum ΔF/F value was used for detection threshold.

### Quantitative PCR (qPCR)

Total RNA was isolated using the Direct-zol RNA Kits according to the manufacturer’s instructions. Reverse transcription was performed using ProtoScript II Reverse Transcriptase with a mixture of oligo(dT) and random primers to generate cDNA. qPCR was performed using a two-step amplification protocol on an Agilent Mx3005P real-time PCR system. Reactions were prepared using Maxima SYBR Green (Thermo). Relative gene expression levels were calculated using the ΔΔCt method. Primers (sequence 5′→3′): SV40-LT: TAAAGCATTGCCTGGAACGC (forward), AAACTCAGCCACAGGTCTGTAC (reverse); Map1a: GGAGACCTTATCCTACAGAGTGG (forward), CCCTAAGCAGGAAACAGTGAGG (reverse); Fos: CGGGTTTCAACGCCGACTA (forward), TTGGCACTAGAGACGGACAGA (reverse); Srrm2: TGGTCCAGGTCCTCGGATTC (forward), TGTCAAGGCAAGTGCAATTCT (reverse).

### Ribosome pulldown and RNA sequencing

Ribosome pulldown was based on the previous study^15^ with modifications. Neurons were lysed with lysis buffer (20 mM Tris pH 7.4, 140 mM KCl, 5 mM MgCl_2_, 1 mM DTT, 1% Triton X-100, EDTA-free protease inhibitor (Roche cOmplete), SUPERase-In (Thermo), Murine RNase inhibitor (NEB)). Lysates after centrifugation was subjected to free biotin removal using Zeba 7k spin columns equilibrated with low-salt buffer (the same to lysis buffer with Triton X-100 adjusted to 0.1%), following manufacturer’s guideline. Biotinylated ribosomes were enriched using 100 μl of MyOne streptavidin C1 magnetic Dynabeads (Invitrogen) per approx. 1 × 10^6^ cells. The beads washed twice with Buffer A (100 mM NaOH, 50 mM NaCl), once with Buffer B (100 mM NaCl) and once with low-salt buffer before binding. Ribosome enrichment was done via overnight incubation with rotation at 4 °C. The next day, the beads were washed three times for 20 min at 4 °C with roration using high-salt wash buffer (20 mM Tris pH 7.4, 500 mM KCl, 5 mM MgCl2, 1 mM DTT, 1% Triton X-100, EDTA-free protease inhibitor (Roche cOmplete), SUPERase-In (Thermo), Murine RNase inhibitor (NEB)). After high-salt wash, the beads were moved into a new tube with the low-salt buffer, supernatant was removed, and RNA extraction was performed using TRIzol and followed by RNA precipitation aided by GlycoBlue (Invitrogen). RNA quality control and concentration measurement was performed using Agilent 4200 TapeStation and Qubit (Invitrogen). RNA-seq libraries were generated using NEBnext Ultra II Directional Library Prep Kit (NEB).

Paired-end RNA-seq (2 × 100 bp) was generated using the Illumina NextSeq 2000. Gene ontology and network analysis was performed with R (4.5.1) package clusterProfiler (4.17.0)^50^. Plots were generated with ggplot2 (3.5.2)^49^.

### scRNA-seq analysis

scRNA-seq results from DIV23 mouse primary cortical neurons were provided by Krizay D. and Boland MJ^24^. Raw count data were subset to 2000 highly variable genes using scanpy’s ‘pp.highly_variable_genes’ function (flavor="seurat_v3"). These counts were input to scVI (run with default settings) to get low dimensional representation of the data. scanpy was used to visualize data: the ‘pp.neighbors’ function was used to get cell nearest neighbors based on scVI latent representations, then ‘tl.umap’ (min_dist=0.3) to get cell embeddings in 2D, and finally data were plotted with ‘pl.umap’. Finally, cell types were annotated based on manual inspection of cell type marker gene expression. Python 3.12.3, anndata 0.10.7, scvi-tools 1.1.2, scanpy 1.10.1 were used.

### Alternative splicing analysis

rMATS v4.3.0 was run (readLength=50) with samples from two different conditions. The output was then filtered for splicing events with P-adjusted < 1e-05 and a minimum delta PSI of 0.1. The results were plotted with the ggplot2 (v4.0.0) R package.

### Splicing isoform visualization

To generate cDNA from pulldown or total RNA samples, reverse transcription was performed using ProtoScript II Reverse Transcriptase with a mixture of oligo(dT) and random primers to generate cDNA. Splicing isoforms of *Anks1b* and *Cd47* were amplified with Q5 polymerase (NEB) and visualized using agarose gel 3.3 - 3.5%. Anks1b: 58 °C annealing, 45 sec elogation. Primers (sequence 5′→3′): tccccgatcaaagctggagag (forward) tttccggatgatgctgccagtac (reverse). Cd47: 56 °C annealing, 1 min elongation. Primers: ggtttggggatcatagctctagca (forward), ccttccagctgtgagtcgtgaag (reverse).

## Data Availability

The data generated in this study have been deposited in the NCBI Gene Expression Omnibus (GEO) under accession number GSE333165. The dataset is currently private and available to editors and reviewers through the GEO reviewer access link/token provided during manuscript submission. The data will be released publicly upon publication of this article.

## Code Availability

The code used in this study is available from the corresponding author upon request and will be released publicly upon publication of this article.

## Acknowledgments

We are grateful to Bateup H, Aisenberg E, Gomez A, and Hsiao M, and the members of the Ingolia and Lareau labs for discussion. We appreciate Krizay D. and Boland M. for sharing scRNA-seq data. S.L. appreciates the Korea–U.S. Special Exchange Program for STEM Students (KORUS) and Simms E. Plasmids were provided by the Weissman and Ullman labs via addgene. Imaging was performed at UC Berkeley Molecular Imaging Center (RRID:SCR_017852).

## Author Contributions

J.W.K. and N.T.I. conceived the study. J.W.K. performed all experiments unless otherwise stated, with help from J.H.A.Y. and S.L. R.E. designed and performed alternative splicing and scRNA-seq data analyses. N.T.I., L.F.L., T.H., and Y.N.J. provided resources and scientific guidance. J.W.K. and N.T.I. wrote the manuscript with input from all authors.

---

**Extended Data Fig. 1.**
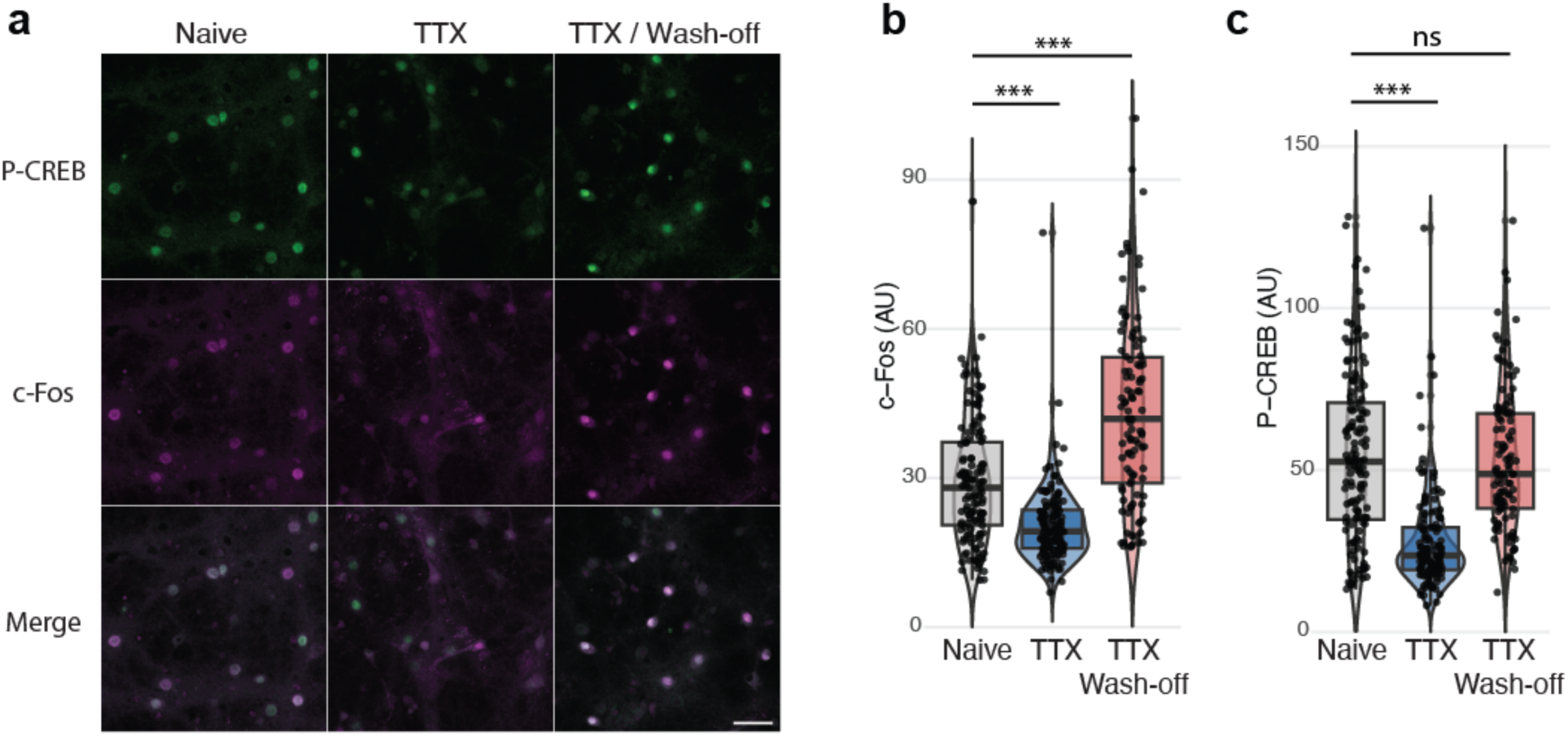
Endogenous c-Fos protein levels and network activity. (**a** to **c**) Endogenous c-Fos protein levels with immunocytochemistry. TTX (1 µM) was treated for 2 hours, primary mouse cortical neurons (DIV 23) were fixed 2 hours after wash-off. Statistical analysis: one-way ANOVA with Tukey’s post-hoc test. (b) n = 382, F = 69.98, p < 2e-16 (Naive vs. TTX), p < 2e-16 (Naive vs. TTX wash-off). (c) n = 378, F = 61.59, p < 2e-16 (Naive vs. TTX), p =0.9713 (Naive vs. TTX wash-off). *** p < 0.001. Scale bars: 50 µm.

**Extended Data Fig. 2.**
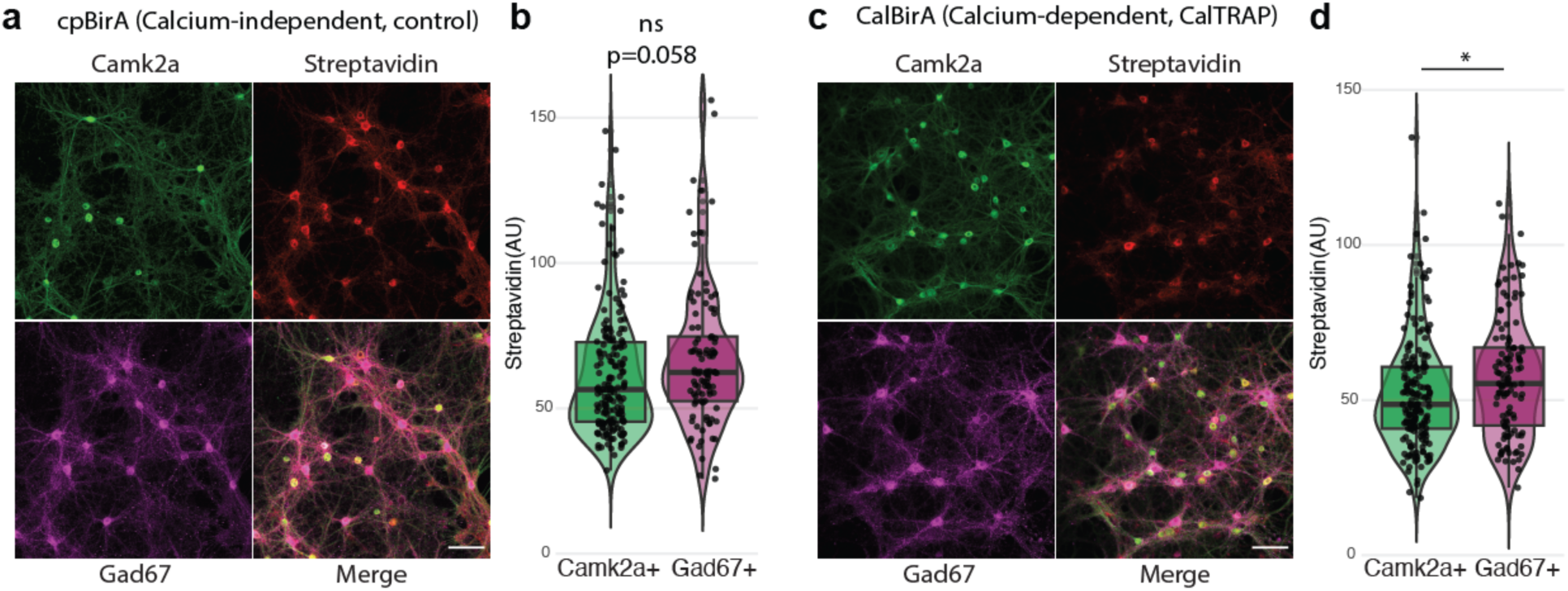
CalTRAP labeling and neuronal cell types. (**a** to **d**) CalTRAP biotinylation levels are compared between excitatory and inhibitory neurons using immunocytochemistry. Primary cortical neurons at DIV 23. (a and b) Calcium-independent labeling with cpBirA. (c and d) Calcium-dependent labeling with CalBirA. Statistical analysis: two-way ANOVA. cpBirA vs. CalBirA: F = 30.432, p = 5.25e-08; Camk2a+ vs. Gad67+: F = 9.295, p = 0.0024; interaction: p = 0.9858. (b) n = 175 (Camk2a+), n = 101 (Gad67+), median: 56.46 (Camk2a+), 62.384 (Gad67+). Tukey’s post-hoc test: p = 0.058. (d) n = 181 (Camk2a+), n = 117 (Gad67+), median: 48.618 (Camk2a+), 55.355 (Gad67+). Tukey’s post-hoc test: p = 0.0143. * p < 0.05. Scale bars: 50 µm.

**Extended Data Fig. 3.**
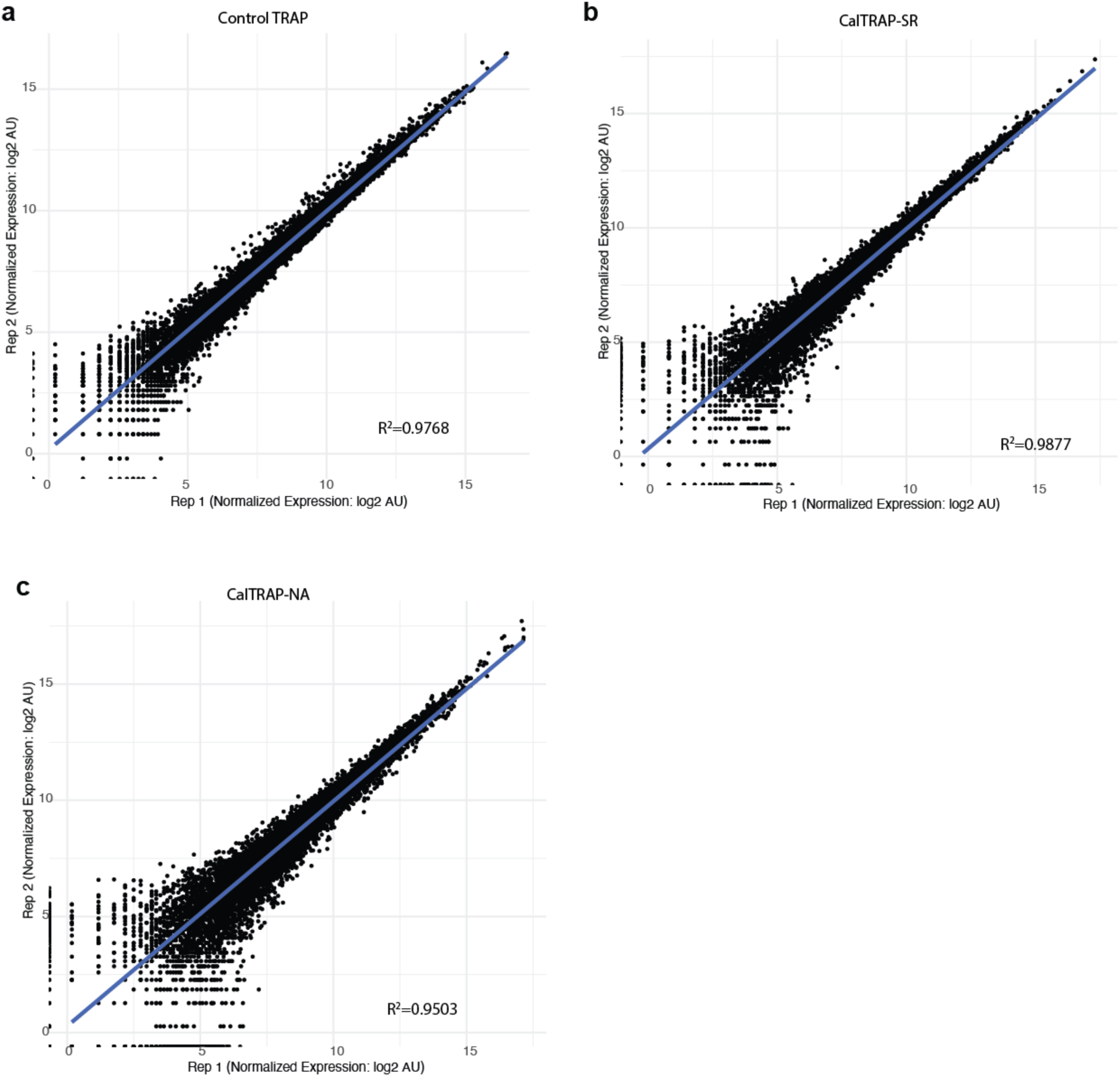
Reproducibility of CalTRAP-seq results. (**a** to **c**) Comparing normalized expression abundances between biological replicates. Total 13781 genes. (a) Control TRAP, R^2^ = 0.9768. (b) CalTRAP-SR, R^2^ = 0.9877. (c) CalTRAP-NA, R^2^ = 0.9503.

**Extended Data Fig. 4.**
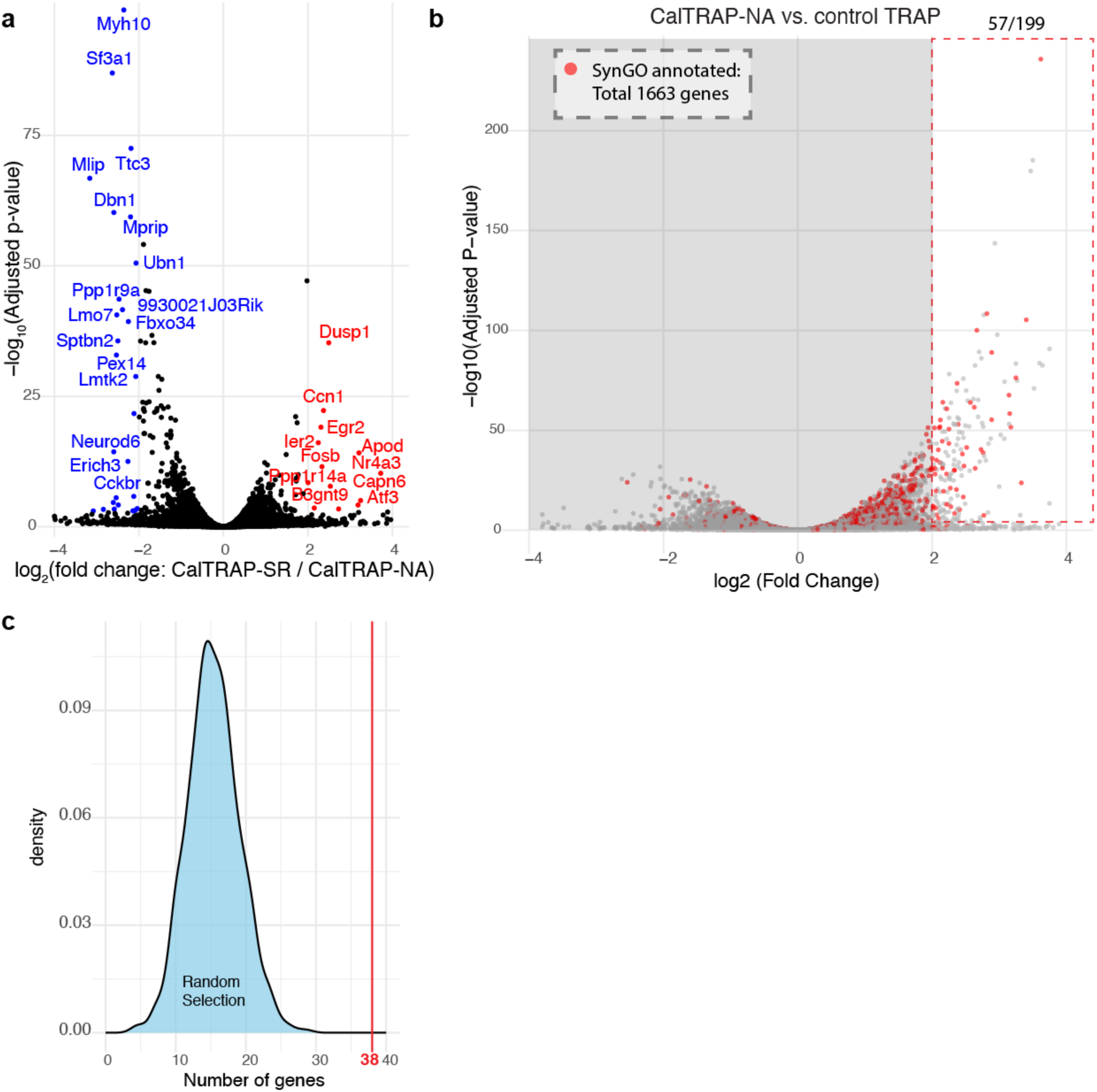
Analyses of gene expression from network activity-engaged neurons. (**a**) CalTRAP-seq differential gene expression analysis comparing CalTRAP-SR vs. CalTRAP-NA. Genes with adjusted p-value < 0.001 and log2FoldChange > 2 or < −2 are highlighted. (**b** and **c**) SynGO and SFARI gene enrichment from top enriched 199 genes from CalTRAP-NA vs. Control TRAP (log2FoldChange > 2, adjusted p-value < 0.001). (c) SynGO genes show higher expression levels in CalTRAP-NA. 57/199 genes from top enriched genes. (d) SFARI: 38 genes (category 1 & 2). Selecting 38 from 199 by chance is p < 1.19187e-09 (distribution from 10,000 random trials).

**Extended Data Fig. 5.**
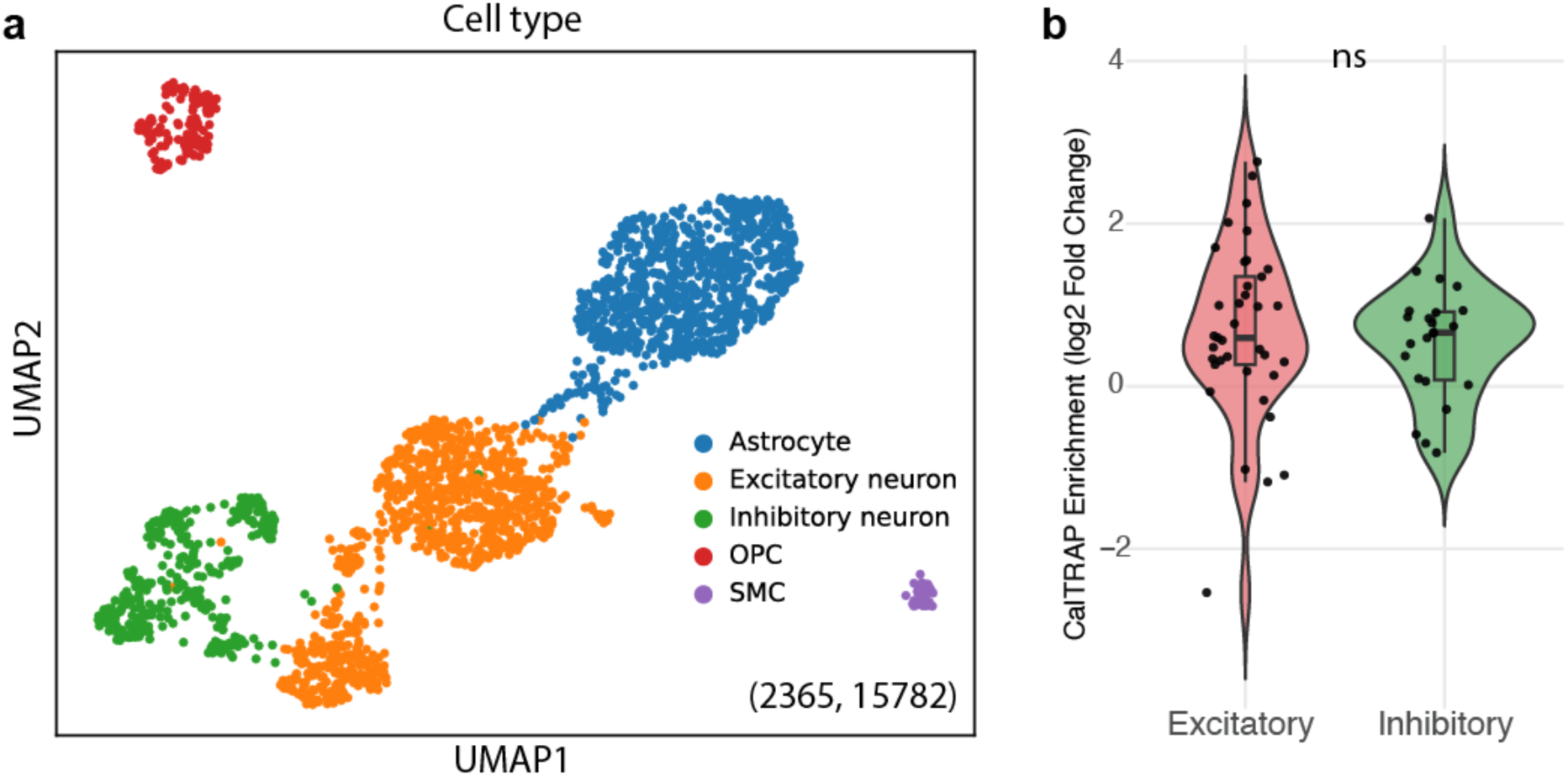
CalTRAP labeling in different neuronal types. (**a**) Cell type clustering of scRNA-seq results from 2356 mouse primary neurons at DIV 23 (Krizay et al. 2022). (**b**) Top differentially expressing genes between excitatory and inhibitory neurons were selected from (a). Excitatory: 37, inhibitory: 23 top gene were selected with log2foldchange > 2.5 and adjusted p-value < 0.001. Their enrichment in CalTRAP-NA (vs. Control TRAP) is shown. Welch’s t-test, p = 0.5981.

**Extended Data Fig. 6.**
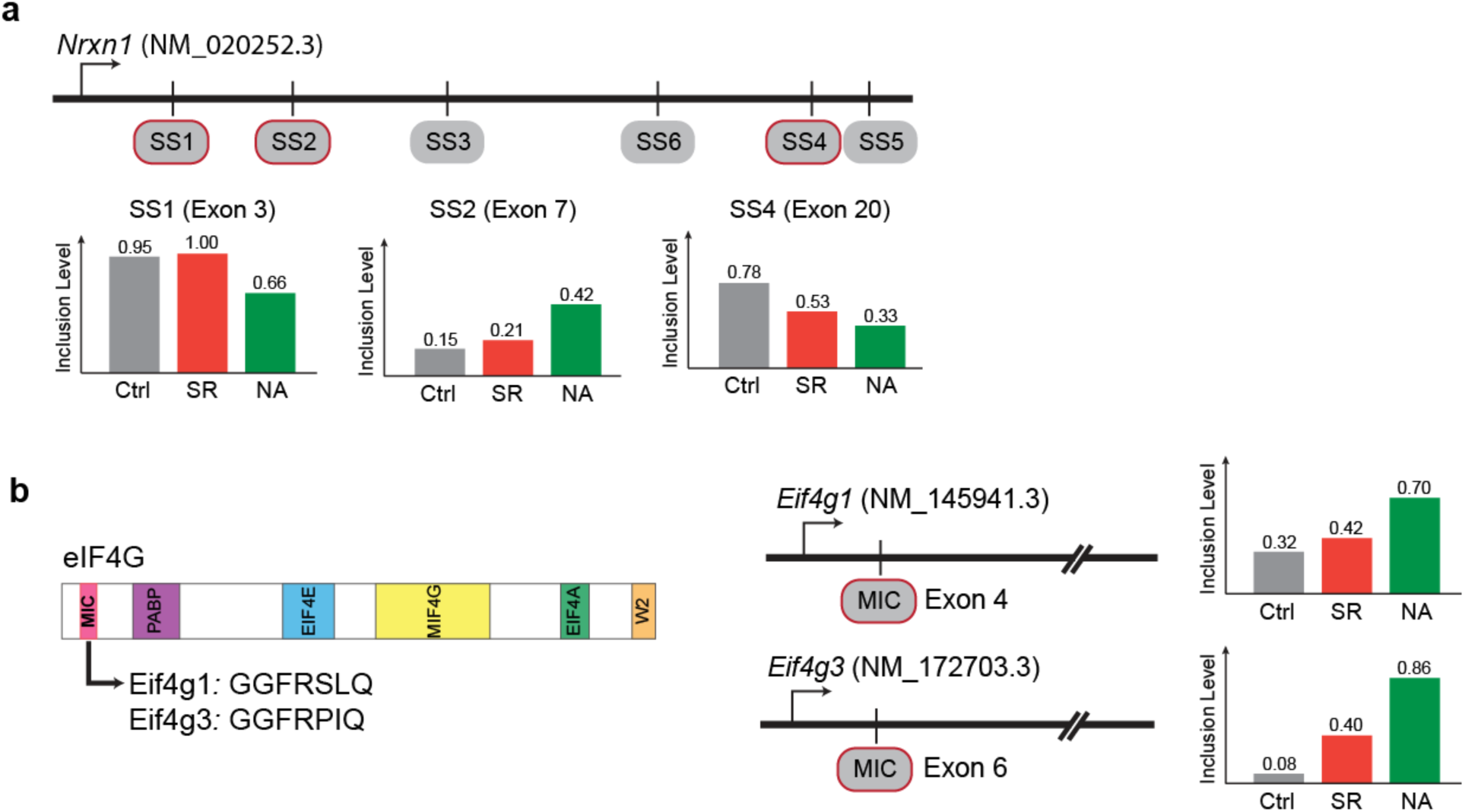
Network activity-dependent alternative splicing of Nrxn1 and Eif4g1/3. (**a**) 3 alternative splicing events from *Nrxn1* gene were identified between CalTRAP-NA and control TRAP. FDR: 6.51e-09 (SS1), 5.44e-12 (SS2), 3.37e-08 (SS3) from rMATS. All three sites don’t show significant differences (FDR < 1e-6) between CalTRAP-SR and control TRAP comparison. (**b**) Microexons in *Eif4g1* and *Eif4g3* genes show network activity-dependent alternative splicing. FDR: 8.66e-15 (Eif4g1), 3.44e-15 (Eif4g3) from rMATS. None shows statistical significance (using FDR < 1e-6) from CalTRAP-SR vs. control TRAP comparison.

**Extended Data Fig. 7.**
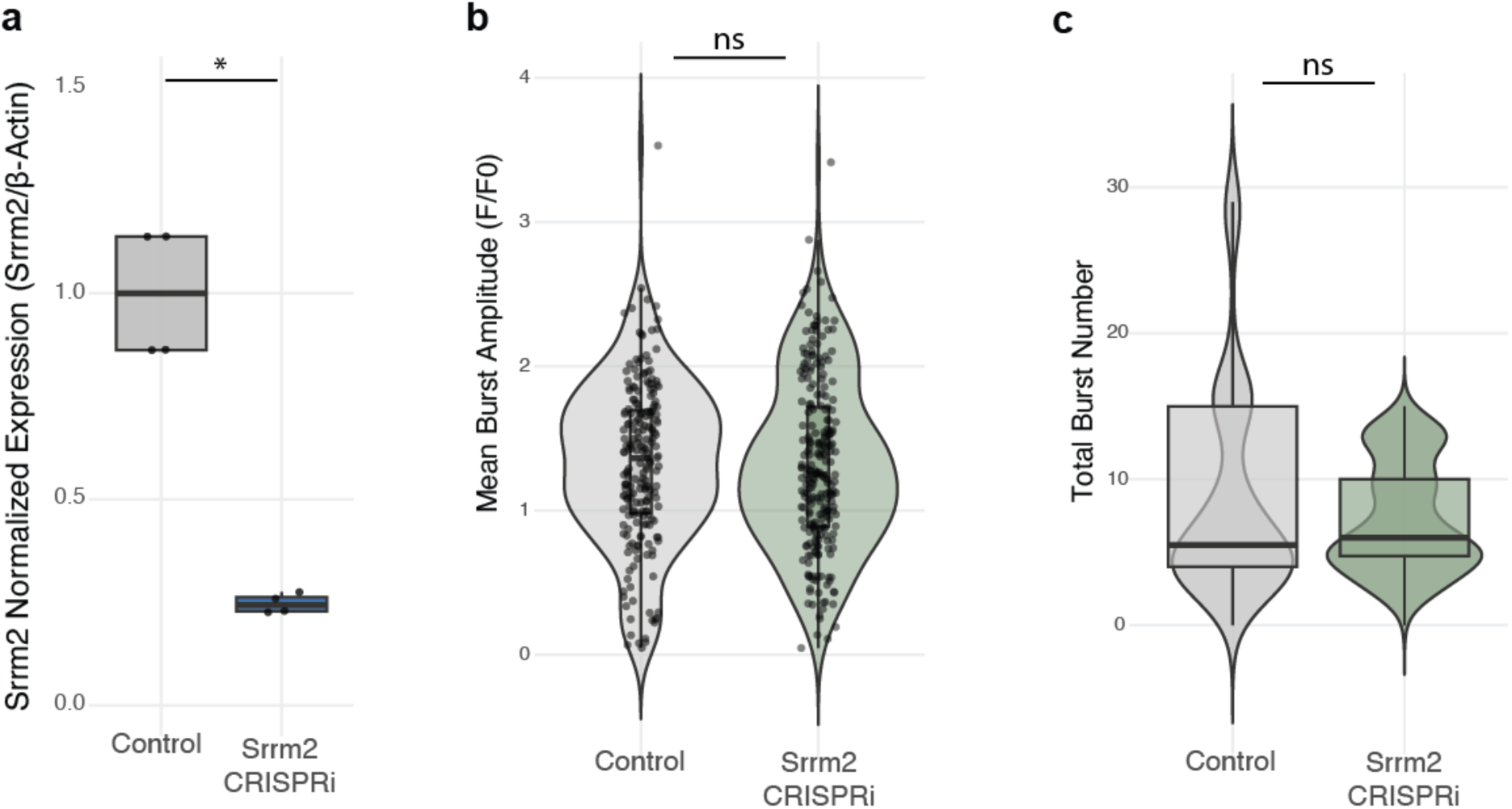
CRISPR interference of Srrm2 in mouse cortical neurons. (**a**) CRISPR interference shows approx. 4-fold reduction of *Srrm2* expression levels. n = 4, Wilcoxson signed-rank test, p = 0.02857. (**b**) Mean burst amplitude from n = 210 from 11 biological replicates (control), n = 251 from 13 biological replicates (CRISPRi). Welch’s t-test, p = 0.804. (**c**) Total spike numbers comparing 11 (control) and 13 (CRISPRi) independent measurements, see Figs. 6b and 6c. Wilcoxson signed-rank test, p = 0.7397.

